# Tissue-based IL-10 signalling in helminth infection limits IFNγ expression and promotes the intestinal Th2 response

**DOI:** 10.1101/2021.08.10.455867

**Authors:** Holly C. Webster, Virginia Gamino, Amy L. Shergold, Anna T. Andrusaite, Graham A. Heieis, Simon W. F. Milling, Rick M. Maizels, Georgia Perona-Wright

**Author notes:** Corresponding author: Dr Georgia Perona-Wright, Institute of Infection, Immunity and Inflammation, University of Glasgow, 120 University Place, Glasgow, G12 8TA, Soctland, UK.

## Abstract

Type 2 immunity is activated in response to both allergens and helminth infection. It can be detrimental or beneficial, and there is a pressing need to better understand its regulation. The immunosuppressive cytokine IL-10 is known as a T helper 2 (Th2) effector molecule, but it is currently unclear whether IL-10 dampens or promotes Th2 differentiation during infection. Here we show that helminth infection in mice elicits IL-10 expression in both the intestinal lamina propria and the draining mesenteric lymph node, with higher expression in the infected tissue. *In vitro*, exogenous IL-10 enhanced Th2 differentiation in isolated CD4^+^ T cells, increasing expression of GATA3 and production of IL-5 and IL-13. The ability of IL-10 to amplify the Th2 response coincided with its suppression of IFNγ expression and, *in vivo*, we found that, in intestinal helminth infection, IL-10 receptor expression was higher on Th1 cells in the small intestine than on Th2 cells in the same tissue, or on any Th cell in the draining lymph node. *In vivo* blockade of IL-10 signalling during helminth infection resulted in an expansion of IFNγ^+^ and Tbet^+^ Th1 cells in the small intestine and caused a coincident decrease in IL-13, IL-5 and GATA3 expression by intestinal T cells. Together our data indicate that IL-10 signalling promotes Th2 differentiation during helminth infection at least in part by regulating competing Th1 cells in the infected tissue.

## 1 Introduction

Gastrointestinal helminths infect more than 1.5 billion people per year^1^ and type 2 immune responses are critical for parasite expulsion^2, 3^ and subsequent wound healing^4^. The same type 2 responses can be harmful in contexts such as allergy and atopic asthma^5^. Better understanding of type 2 immunity is important both to optimise anti-helminthic strategies, such as vaccination, and to accelerate new therapeutic approaches to atopic diseases. Type 2 immunity is initiated when antigen or allergen exposure coincides with the release of alarmins such as interleukin (IL) −25, IL-33 or thymic stromal lymphopoietin (TSLP)^6^. Alarmins promote the activation of type 2 innate lymphoid cells (ILC2) and the recruitment and activation of dendritic cells (DCs) that direct T helper 2 (Th2) cell differentiation^7–9^. Th2 cells secrete cytokines such as IL-4, IL-5 and IL-13, which drive further Th2 polarisation, direct B cell class-switching, recruit effector cells such as eosinophils, basophils and mast cells, and stimulate goblet cell hyperplasia, mucus secretion, epithelial turnover and increased smooth muscle reactivity^10, 11^. Th2 cytokines show spatial patterning, with IL-4 concentrated in the lymph node and IL-5 and IL-13 in the effector tissues^12–15^, reflecting the timing of their production, and their distinct target cells^16^. If cytokine production becomes chronic or excessive, type 2 immunity can drive fibrosis, scar formation and loss of tissue function. Regulatory mechanisms are therefore an inherent part of type 2 immunity, balancing protective immunity and immunopathology.

IL-10 is a key regulatory cytokine. It was first described as a Th2 effector cytokine, secreted by isolated Th2 clones^17, 18^. Neonates have a Th2 bias that correlates with high expression of IL-10^19, 20^. DC-derived IL-10 has been reported to promote antigen-specific Th2 responses in a model of allergic dermatitis^21^, and IL-10-dependent induction of STAT3 and Blimp-1 has recently been shown to be essential for the development of inflammatory Th2 cells in the lung during asthma^22, 23^. IL-10 has also been reported to support antibody isotype switching^24^ and to amplify mast cell activity^25–27^.

However, IL-10 is known foremost as a suppressive cytokine, particularly in Th1 responses. IL-10-deficient mice develop spontaneous colitis^28^, driven by exaggerated IFNγ and IL-17 responses to commensal bacteria^29–31^. Protozoan, viral and bacterial infections in IL-10-deficient mice also show potent increases in Th1 and Th17 cytokines, associated with rapid pathogen clearance but also with acute and often fatal immunopathology^32–34^. The original description of IL-10 as a Th2 effector molecule *in vitro* may reflect its ability to limit Th1 differentiation, especially *in vitro* where Th1 and Th2 responses are mutually antagonistic^18^. The impact of IL-10 on Th2 responses *in vivo*, in the context of mixed T cell responses, is less clear.

IL-10 expression increases in the draining lymph node during infection with the murine helminth *Heligmosomoides polygyrus*^35–37^, and it is essential for host survival during infection with the whipworm *Trichuris muris*^38^. The Th2 response to *Nippostrongylus brasiliensis* infection also requires IL-4 dependent IL-10 signalling^39^. IL-10 has different effects at different stages of *Trichinella spiralis* infection, promoting intestinal mast cell accumulation and clearance of adult worms, but suppressing the immune response against larvae encysted in peripheral muscle^25^. The tissue location of IL-10 activity may be influential. The impact of IL-10 is dependent on timing, location and cell type^34, 40^ and yet the cells that IL-10 targets at the site of helminth infection, and the intestinal networks by which it acts, are still uncertain. Previous studies of T cell regulation during helminth infection have focused on lymph node responses, limited by the technical difficulties created by the extensive mucus production, oedema, and tissue fragility in the helminth-infected gut. We and others recently published new protocols for successful isolation of intestinal immune cells during active type 2 immune responses^40–42^. Here, we have used these technical advances to interrogate the impact of IL-10 on the regulation of the Th2 immune response in the infected tissue during an enteric helminth infection.

Our data show that, during infection with the intestinal helminth parasite *H. polygyrus*, IL-10 is a striking feature of the immune response in the infected intestinal tissue. We demonstrate *in vitro* that IL-10 promotes Th2 cytokine expression in unpolarised cells in part by suppressing IFNγ expression. We show in vivo that *H. polygyrus* infection includes an intestinal Th1 response that is limited by direct IL-10 signalling. Surface expression of the IL-10 receptor was higher on Th1 cells in the infected small intestine than on Th2 cells locally or in the draining lymph node, and in vivo blockade of IL-10 signalling during *H. polygyrus* infection resulted in enhanced Th1 and reduced Th2 activity in the small intestine. Together our data suggest a regulatory loop in helminth-infected, intestinal tissue in which IL-10 suppresses competing Th1 cells to promote Th2 immunity.

## 2 Results

### IL-10 expression increases in H. polygyrus infection and is higher in the small intestine than the draining lymph node

To investigate the impact of IL-10 on the immune response to intestinal helminth infection, we first assessed the location of IL-10 expression during infection with the enteric roundworm, *Heligmosomoides polygyrus*. IL-10 production has previously been shown to increase during *H. polygyrus* infection in cells of the draining, mesenteric lymph node (MLN)^35–37, 43^. Using IL-10 reporter mice^44, 45^, we saw a significant increase in the percentage of CD45^+^ IL-10^+^ cells upon infection in the MLN (**Figure 1A**) but also in the small intestine lamina propria (SILP). Expression was significantly higher in the small intestine compared with MLN (**Figure 1A**). We then analysed the cells producing IL-10 in both the SILP and MLN. Our gating strategies are shown in figures S1 and S2. In the SILP, the percentage of CD8^+^ T cells, B cells and ILCs expressing IL-10 increased at day 7 (D7) post-infection with *H. polygyrus* compared to naïve controls, whereas the proportion of IL-10^+^ CD4^+^ T cells remained unchanged and there was a small decrease in IL-10^+^ myeloid cells (**Figure 1A–F**). In the MLN, the percentage of IL-10^+^ CD8^+^ T cells, CD4^+^ T cells and ILCs increased upon infection, while both IL-10^+^ B cells and myeloid cells remained unchanged between naïve and infected samples (**Figure 1D & 1F**). IL-10 expression in both naïve and infected mice was higher in the SILP compared to the MLN In all cell types (**Figure 1A–F**). We then compared the contribution of each cell type to the total IL-10^+^ population (**Figure 1G–H**). In the SILP, CD4^+^ T cells and myeloid cells made up the majority of CD45^+^ IL-10^+^ cells in both naïve and infected mice. The proportion of IL-10^+^ cells that were CD8^+^ T cells also expanded in the SILP during infection (**Figure 1G**). In the MLN, CD4^+^ T cells were the dominant population of CD45^+^ IL-10^+^ cells in both naïve and infected animals, although the proportion of B cells and ILCs within the IL-10^+^ pool increased slightly upon infection (**Figure 1H**). Together, these data show multiple sources of IL-10 active during *H. polygyrus* infection and demonstrate high IL-10 expression at the site of infection, in the SILP.

**Figure 1.**
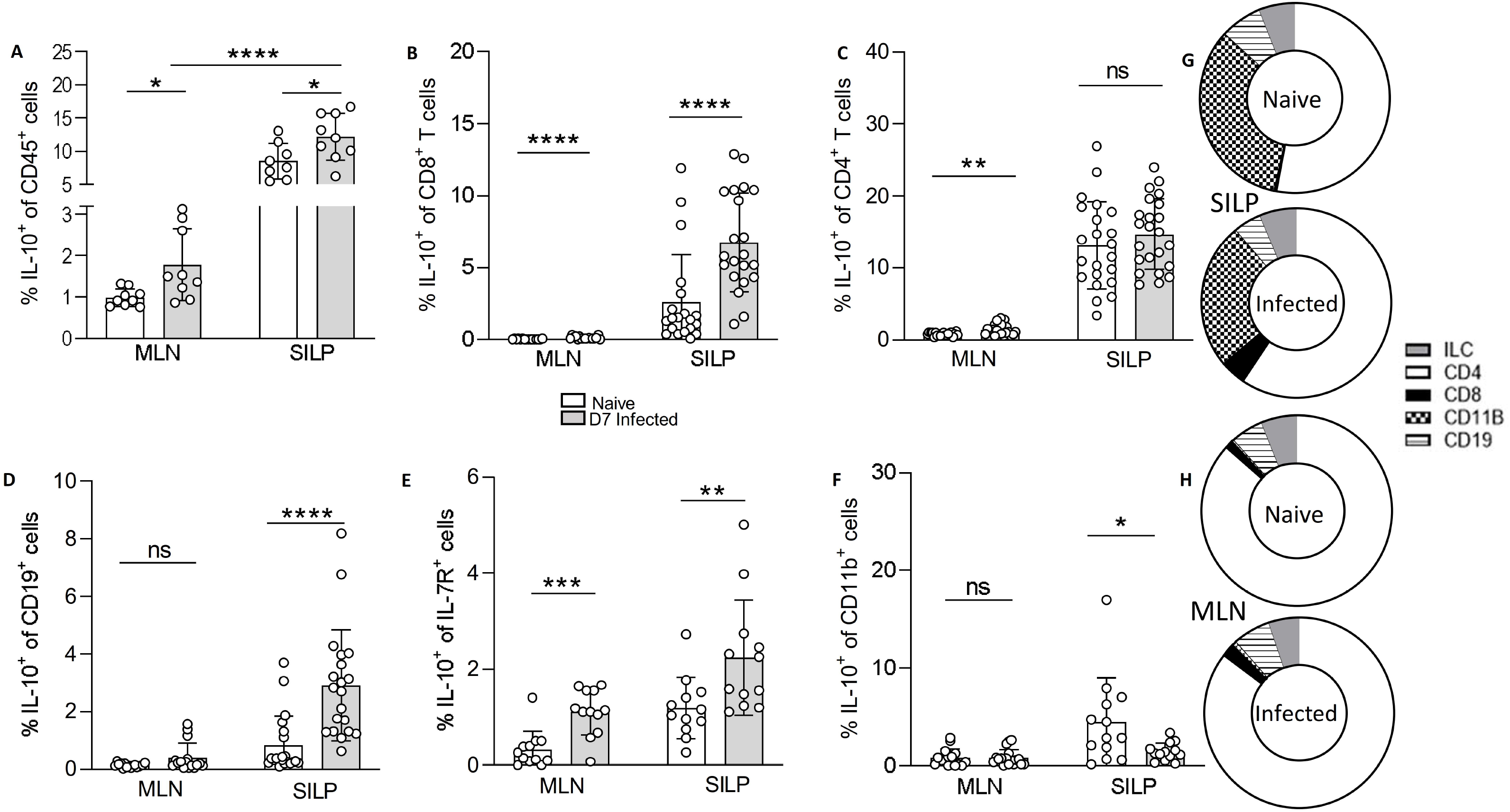
IL-10 expression increases in the MLN and small intestine during *H. polygyrus* infection. Il10gfp-foxp3rfp B6 mice were infected with 200 L3 *H. polygyrus* and 7 days later the small intestine and MLN collect for analysis. % of IL-10^+^ of (A) CD45^+^, (B) CD8^+^, (C) CD4^+^, (D) CD19^+^, (E) IL-7R^+^ and (F) CDllb^+^ cells from the MLN and small intestine of na“ive and D7 infected mice. Changes in proportion of IL-10 producing cells from total co45^+^ IL-10^+^ cells in the (G) small intestine and (H) MLN from naïve and D7 infected mice Graphed data are shown with mean ± 1 SD and are pooled from 3 independent experiments with *n*=2-5 per experiment. Statistical significance was calculated by Student *t* test where data were normally distributed (A, C (SILP)) and Mann Whitney U test where data were not normally distributed (B, D (MLN), D, E, F). (Significance \**p*<⍰0.05, \*\**p*<⍰0.01, \*\*\**p*<⍰0.001, \*\*\*\**p*<⍰.0001).

### IL-10 enhances Th2 differentiation in vitro

To determine the functional importance of IL-10 expression during *H. polygyrus* infection, we first considered whether direct IL-10 signalling to CD4^+^ T cells could contribute to Th2 polarisation. We stimulated purified CD4^+^ T cells with αCD3, αCD28 and IL-2 *in vitro* (Th0 cultures) (Figure S3) and added recombinant IL-10. The presence of IL-10 caused a significant increase in the expression of GATA3, IL-5 and IL-13 (**Figure 2A & B**), showing that, in an unpolarised CD4^+^ T cell, IL-10 can enhance Th2 differentiation. When we added IL-10 to polarised Th2 cell cultures (CD4^+^ T cells stimulated with αCD3, αCD28, IL-2, IL-4 and anti-IFNγ), the presence of IL-10 did not result in a further significant increase in expression of GATA3 or of the effector cytokines IL-5 and IL-13 (**Figure 2C & D**), perhaps reflecting the high levels of these cytokines already produced under polarising conditions. The impact of IL-10 in enhancing Th2 differentiation in Th0 cultures appeared to be on polarisation rather than on activation or proliferation, since neither CD44, CD69 nor cell division were significantly different in Th0 cells cultured in the presence or absence of IL-10 (**Figure 2E, F, G**). Together, these data show that IL-10 can act directly on CD4^+^ T cells to promote Th2 polarisation, particularly in sub-maximal polarisation conditions, and that this occurs independently of activation and proliferation.

**Figure 2.**
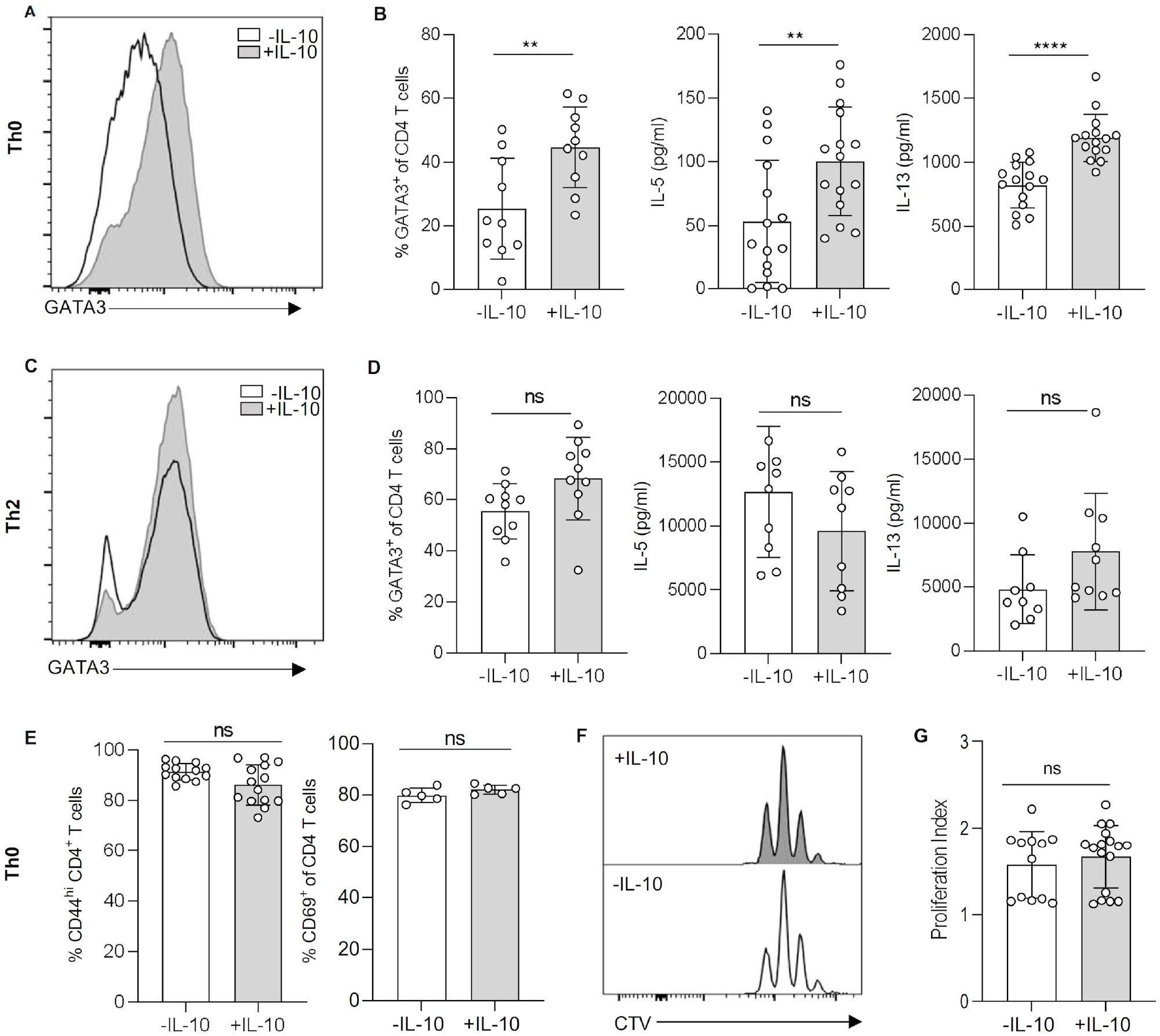
IL-10 enhances Th2 differentiation independently of activation and proliferation. *In vitro* polarised ThO and Th2 cells were cultured with aCD3, aCD28 and IL-2 for 4 days with or without IL-10. (A) Representative histogram of GATA3 staining from IL-10 stimulated and unstimulated ThO cells. (B) % of CD4^+^ GATA3^+^ CD4 T cells (left) and pg/ml of IL-5 (middle) and IL-13 (right) in ThO culture supernatants. (C) Representative histogram of GATA3 staining from IL-10 stimulated and unstimulated ThO cells. (D) % of CD4^+^ GATA3^+^ CD4 T cells (left) and pg/ml of IL-5 (middle) and IL-13 (right) in Th2 culture supernatants. (E) % of CD44^hi^ (left) and CD69^+^ CD4 T cells from IL-10 stimulated and unstimulated ThO cultures. (F) Representative histograms of cell trace violet staining from IL-10 stimulated (right) and unstimulated (left) ThO cultures. Graphed data are shown with mean ± 1 SD and are pooled from 3 independent experiments with *n*=4-5 per experiment. Statistical significance was calculated by Student *t* test where data were normally distributed (B, D, E) and Mann Whitney U test where data were not normally distributed (G). (Significance \*\**p*<⍰ 0.01, \*\*\*\**p*<⍰.0001).

### Th2 induction by IL-10 *in vitro* coincides with suppression of IFNγ

An inverse relationship between IL-10 and IFNγ has been well described^18, 33, 46–48^, and we next aimed to determine if the Th2 skewing effects of IL-10 could be due to IFNγ-mediated suppression. The addition of exogenous IL-10 decreased IFNγ expression in the unpolarised Th0 cells, as well as in polarised Th1 cultures (purified CD4^+^ T cells stimulated with αCD3, αCD28 IL-2 and IL-12) (**Figure 3A**). We then treated Th0 cells with or without IL-10 while blocking IFNγ signalling, using an anti-IFNγ antibody, to test whether the absence of IFNγ signalling could replicate the Th2 polarisation induced by IL-10 alone. Indeed, Th0 cells cultured without IL-10 but in the absence of IFNγ signalling showed equivalent Th2 polarisation to Th0 cells stimulated with IL-10 alone (**Figure 3B**). However, dual treatment of Th0 cells with IL-10 and anti-IFNγ was synergistic and induced higher secretion of IL-5 and IL-13 than each intervention alone (**Figure 3B**). Interestingly, the partial reduction in IFNγ production seen even in strongly polarised Th1 cultures (**Figure 3A**) also corresponded with a rebound in IL-13 expression (**Figure 3C**), although any increase in IL-5 expression did not reach statistical significance (**Figure 3C**). Together, these data suggest that the ability of IL-10 to enhance Th2 polarisation correlates with its ability to limit IFNγ expression.

**Figure 3.**
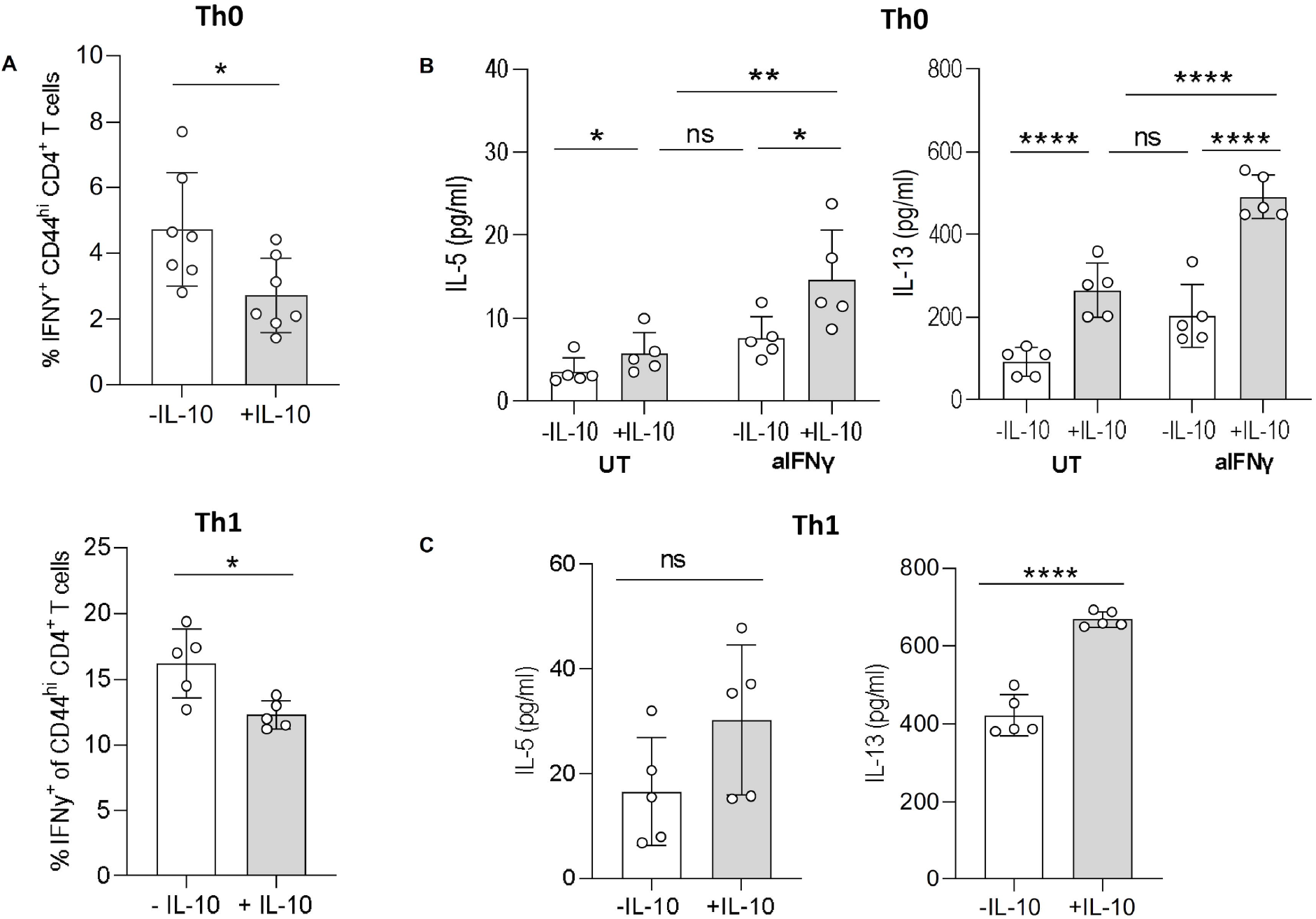
IL-10 suppresses IFNγ expression when promoting Th2 differentiation. *In vitro* polarised Th0 and Thl cells were cultured with αCD3, αCD28 and IL-2 only for Th0, with the addition of IL-12 for Th1 for 4 days with or without IL-10. (A) The% of IFNγ^+^ CD44^hi^ CD4^+^ T cells measured in Th0 (top) and Thl (bottom) cultures. (B) *In vitro* polarised Th0 cells were cultured with αCD3, αCD28 and IL-2 for 4 days with or without IL-10 and with or without anti-lFNy, the concentration of IL-5 and IL-13 measured in supernatants. (C) *In vitro polarised* Thl cells were cultured with αCD3, αCD28 and IL-12 for 4 days with or without IL-10 and IL-5 and IL-13 concentration in the supernatants measured. Graphed data are shown with mean ± 1 SD and are representee of 2 independent experiments with n=S per experiment. Statistical significance was calculated by Student *t* test and one-way ANOVA with Tukey’s post-test for multiple comparisons between groups {Significance *p<0.05, \*\**p*<⍰ 0.01, \*\*\*\**p*<⍰.0001).

### In the infected intestine, IL-10 receptor expression is higher on Th1 cells than Th2

Our *in vitro* data suggested that IL-10 may promote the Th2 response in part by suppressing IFNγ. To compare the potential IL-10 responsiveness of Th1 and Th2 cells *in vivo*, we first measured the expression of IL-10R1, which has been shown to correlate closely with changes in cellular responsiveness to IL-10^48, 49^. We infected B6.4get IL-4 reporter mice^50^ with *H. polygyrus* and identified Th cell subsets using CXCR3 as a marker of Th1 cells and IL-4-GFP as an indicator of Th2 cells. Our gating and isotype controls are shown in supplementary figures S4-S6. As expected in a helminth infection, the frequency and number of IL-4^+^ CD4^+^ Th2 cells increased in both the MLN and small intestine upon *H. polygyrus* infection (**Figure 4A & B**). We were able to define a small population of CXCR3^+^ CD4^+^ Th1 cells in the MLN and SILP (**Figure 4A & B**), the frequency of which decreased in both the MLN and small intestine as the Th2 cells expanded (**Figure 4A & B**). Both CXCR3^+^ Th1 cells and IL-4/GFP^+^ Th2 cells appeared activated, as indicated by scatter profile (**Figure 4C**) and expression of CD44 (**Figure 4D**). CXCR3^+^ CD4^+^ Th1 cells in the small intestine showed significantly higher expression of IL-10R1 than Th2 cells, both in frequency and intensity, in infected and uninfected animals (**Figure 4E–G**). IL-10R1 expression was significantly higher on Th1 cells in the intestine than in the draining MLN, while the lower level of IL-10R1 expression on Th2 cells was similar in both tissues (**Figure 4E–G**). These data indicate that high IL-10R expression is a feature of Th1 cells in the small intestinal mucosa, but not in the draining LN, even during intestinal helminth infection.

**Figure 4.**
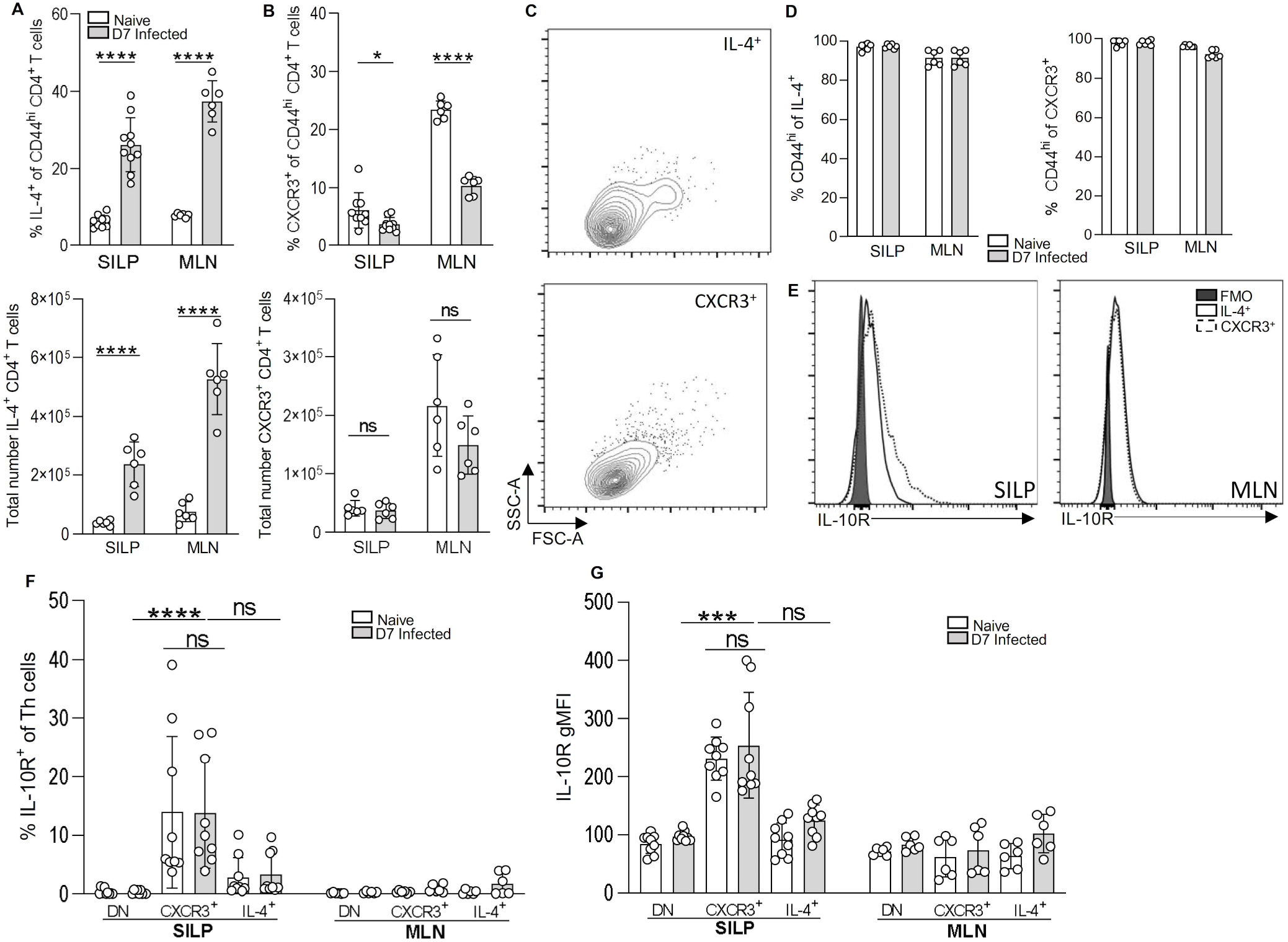
Thl cells in the small intestine during *H. polygyrus* infection show high IL-lOR expression. B6 4get mice were infected with 200 L3 *H. polygyrus* and 7 days post-infection the small intestine and MLN removed. (A) % (top) and total number (bottom) of IL-4(GFP)^+^ Th cells {TCRβ ^+^ CD4^+^ CD44^+^) in the small intestine. (B) % (top) and total number (bottom) of CXCR3^+^ of Th cells in the small intestine. (C) Representative flow cytometry scatter plot of IL-4^+^ (top) and CXCR3^+^ (bottom) Th cells from the SILP of D7 infected mice. (D) % of CD44^hi^ of IL-4(GFP)^+^ (left) and CXCR3^+^ (right) from the SILP and MLN. (E) Representative overlaid histograms of IL-l0R expression of IL-4^+^ and CXCR3^+^ Th cells compared to IL-l0R FMO. (F) Geometric mean of IL-10^+^ expression and (G) % of IL-10^+^ cells of DN (double negative), IL-4(GFP)^+^ and CXCR3^+^ Th cells in the MLN and small intestine. Graphed data are shown with means ± 1 SD and are pooled from 3 independent experiments with n=3 per experiment. Statistical significance was calculated by Student *t* test and Kruskal-Wallis test with Dunn’s post-test for multiple comparisons between groups. (Significance \**p*<0.05 \*\*\**p*<⍰0.001, \*\*\*\**p<*⍰.0001).

### IL-10 signalling blockade in helminth infection leads to Th1 expansion in the infected tissue

Our data so far had shown that IL-10 expression and IL-10 receptor expression were both concentrated at the site of infection and suggested that the primary T cell target of IL-10 signalling during *H. polygyrus* infection may be Th1 cells in the small intestine. To test the impact of IL-10 during infection, we disrupted signalling using a blocking antibody against the IL-10R1. IL-10 blockade during *H. polygyrus* infection caused an increase in the frequency and number of IFNγ^+^ CD4^+^ CD44^hi^ Th1 cells in the intestinal tissue (**Figure 5 A - C**), but no change in the draining MLN (**Figure 5D**). Staining for Tbet^+^ CD4^+^ CD44^hi^ Th1 cells gave the same result (**Figure 5A - D**). These data indicate that, during *H. polygyrus* infection, IL-10 acts to limit Th1 expansion and IFNγ expression in the small intestine.

**Figure 5.**
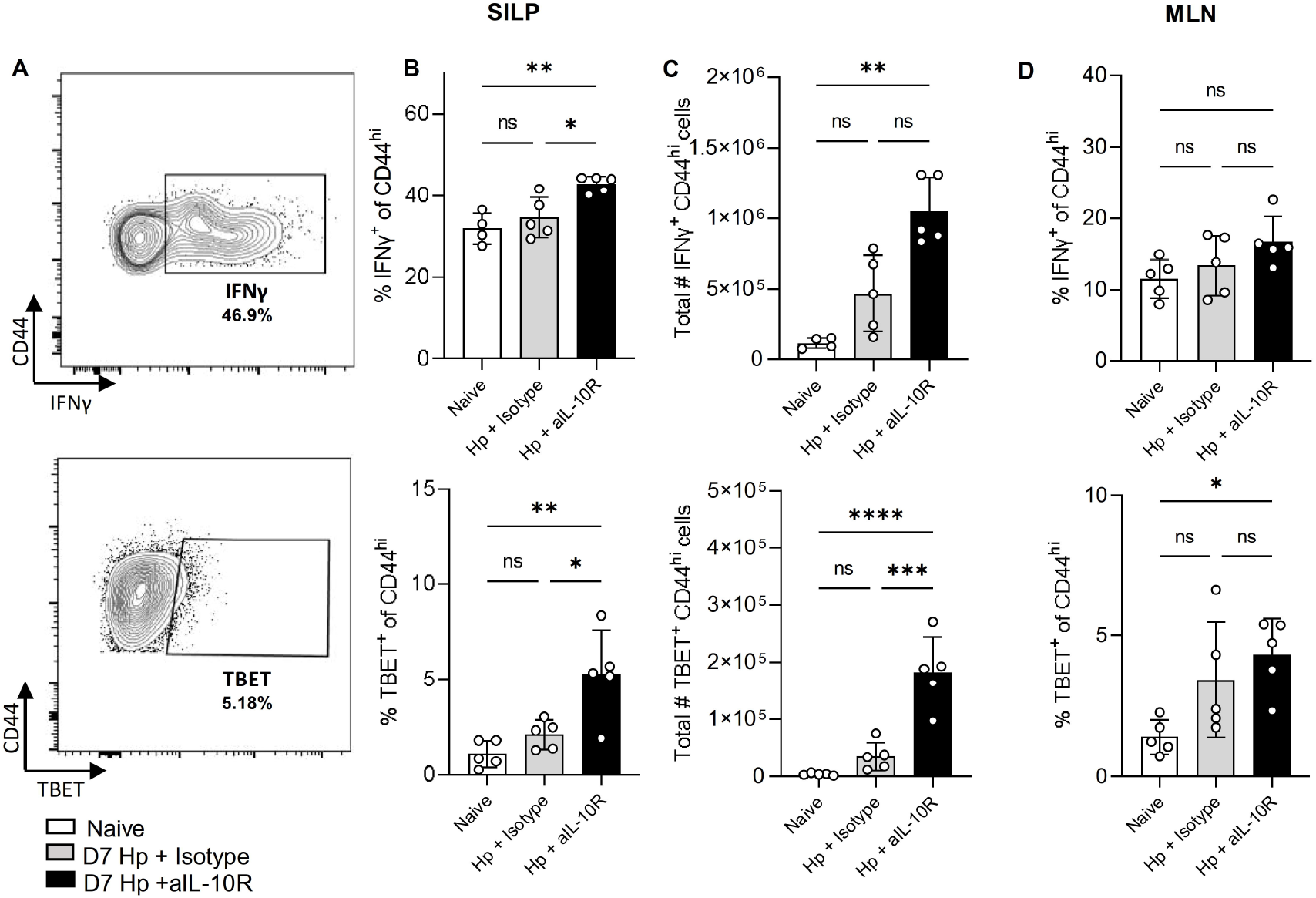
IL-10 signalling blockade in *H. polygyrus* infection leads to expansion of Th1 cells in the small intestine. C57BL/6 mice were infected with 200 L3 *H. polygyrus* and at D-1, D2 and DS treated with anti-IL-10R mAb or isotype control, and 7 days post-infection the small intestine and MLN collect for analysis. Representative flow cytometry of plot IFNγ (top) and TBET (bottom). (B) Percentage of IFNγ+ (top) and TBET^+^ (bottom) of CD44^hi^ CD4^+^ T cells in the SILP. (C) Total number of IFNγ^+^ (top) and TBET^+^ (bottom) of CD44^hi^ CD4^+^ T cells in the SILP. (D) Percentage of IFNγ^+^ (top) and TBET^+^ (bottom) of CD44^hi^ CD4^+^ T cells in the MLN. Graphed data are shown with mean ± 1 SD and are representative of 3 independent experiments with n=4-5 per experiment. Statistical significance was calculated by ANOVA followed by a Tukey’s post-test for multiple comparisons between groups where data were normally distributed (B, C (TBET), D) and Kruskal-Wallis test with Dunn’s post-test for multiple comparisons between groups where data were not normally distributed (C (IFNγ)). (Significance \**p*<0.05, \*\**p*<⍰0.01, \*\*\**p*<⍰0.001, \*\*\*\**p<*⍰.0001).

### Immune competition in the small intestine during H. polygyrus infection is regulated by IL-10

Our observation that IL-10 controls IFNγ expression in the small intestine during *H. polygyrus* infection prompted two hypotheses. The first was that IL-10 signalling might regulate tissue pathology. *H. polygyrus* causes only limited pathology in laboratory mice^51–53^, but parasite larvae cross the intestinal wall twice during infection: by day 2 they penetrate into the sub-mucosa, where they encyst, mature and moult; and again at D7-8 as the adult worms move back into the intestinal lumen, where they persist by twisting themselves around the surface of the host villi. We predicted that, at D7, the effect of the enhanced IFNγ expression in the absence of IL-10 signalling would be to exaggerate pathology around the encysted larvae or at sites of epithelial disruption. In contrast, our data showed that IL-10 blockade had little effect on intestinal pathology at this timepoint. In infected animals, the severity of inflammation was very variable among areas of the tissue sampled, varying from very mild and mostly mucosal at sites distant from the parasite (**Figure 6A, top row**), to severe and submucosal around the encysted parasites (**Figure 6A, bottom left**). Comparing pathology in infected animals treated with IL-10R1 blocking mAb versus those infected but given the isotype control did not reveal any significant differences (**Figure 6A, bottom row**). When we quantified the histology sections, infection was associated with an increase in both combined inflammation and inflammation depth score, but there was no difference in pathology between mice that were infected with or without disrupted IL-10 signalling (**Figure 6B - C**).

**Figure 6.**
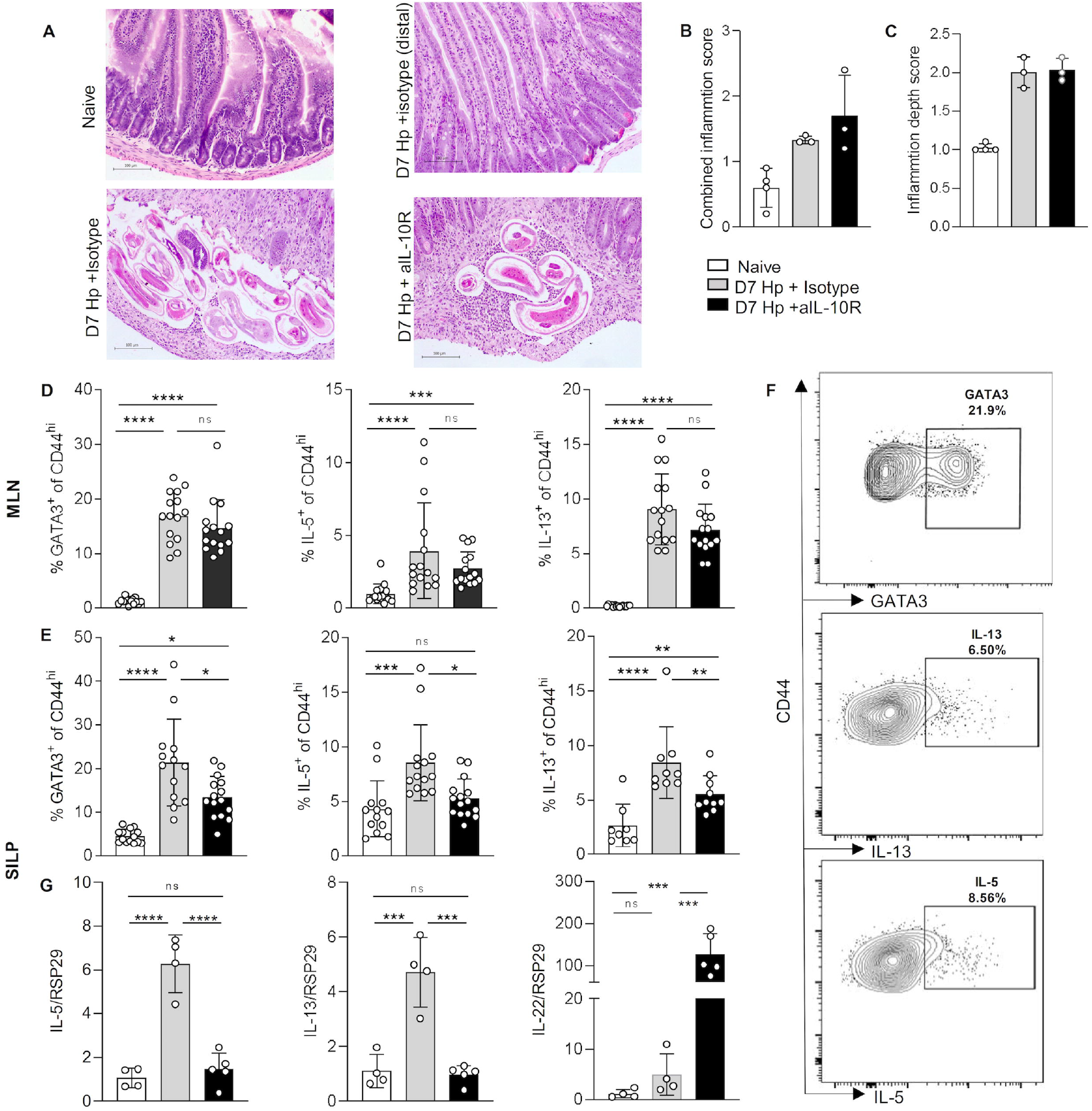
IL-10 signalling promotes the intestinal Th2 response in *H. polygyrus* infection. C57BL/6 mice were infected with 200 L3 *H. po/ygyrus* and at D-1, D2 and DS treated with anti-IL-10R mAb or isotype control, and 7 days post-infection the small intestine and MLN collect for analysis. (A) Representative H&E staining of the duodenum from naïve (top left), D7 isotype parasite area (bottom left), D7 infected distal parasite area mice (top right) and D7 anti-lL-l0R mAb parasite area (bottom right). Histology scoring of (B) inflammation depth and (C) combined inflammation score from the 3 treatment groups. (D) % of GATA3o (left) IL-s+ (middle) IL-13^+^ (right) of CD4^+^ co44^hi^ in the MLN. (E) % of GATA3^+^ (left) IL-5^+^ (middle) IL-13^+^ (right) of CD4^+^ CD44^hi^ in the small intestine. (F) Representative staining of GATA3 (top), IL-13 (middle) and IL-5 (bottom) from *H. polygyrus* infected (anti-IL-10R treated) small intestine. (G) Fold change of IL-5 (left), IL-13 (middle) and IL-22 (right) in the duodenum compared to housekeeping gene (RSP29). Graphed data are shown with mean ± 1 SD and are pooled from 2-3 independent experiments with *n*=4-5 per experiment. Statistical significance was calculated by ANOVA followed by a Tukey’s post-test for multiple comparisons between groups where data were normally distributed (D (IL-5, IL-13), E (GATA3), G) and Kruskal-Wallis test with Dunn’s post-test for multiple comparisons between groups where data were not normally distributed (D (GATA3), E (IL-5, IL-13). (Significance \**p*<0.05, \*\**p*< 0.01, \*\*\**p*<⍰0.001, \*\*\*\**p*<⍰.0001).

Our second hypothesis was that the impact of IL-10 during *H. polygyrus* infection would be to reduce immune competition between Th1 and Th2 cell subsets and thus allow the Th2 response to expand. To assess this, we repeated the infection in the presence or absence of the anti-IL-10R1 blocking antibody and measured the Th2 response. In the MLN, infection induced a clear Th2 response, shown by increased expression of the Th2 master transcription factor GATA3 and the Th2 cytokines IL-5 and IL-13 in CD4^+^ CD44^hi^ cells, and this response was not affected by the blockade of IL-10 signalling (**Figure 6D**). Th2 immunity in the SILP, however, was significantly decreased during IL-10R1 blockade compared with isotype treated controls (**Figure 6E & 6F**). *Il*5 and *Il*13 gene expression in the duodenum, which increased upon infection, was also curtailed by IL-10R1 blockade, becoming similar to levels seen in naïve mice (**Figure 6G**). In contrast, gene expression of *Il*22, a cytokine known to be negatively regulated by IL-10, increased 100-fold during IL-10R1 blockade (**Figure 6G**). Together our data demonstrate that IL-10 signalling limits IFNγ expression by Th1 cells in the small intestine during *H. polygyrus* infection, and that this restriction on IFNγ corresponds with a local expansion of Th2 immune activity in the infected tissue site.

## 3 Discussion

Understanding the regulation of Th2 immunity is important for a variety of diseases, most prominently helminth infection and allergy. IL-10 is a key regulatory cytokine, but while it is well established that IL-10 is suppressive in type 1 immune settings^18^, its role during a type 2 immune response is less well understood. IL-10 expression is known to increase in the lymph nodes and blood during type 2 immune responses^22, 35, 36, 38, 54, 55^, but it has been suggested both to promote and to restrict Th2 immunity^56, 57^. Here we show that, during infection with the helminth *H. polygyrus*, the intestinal immune response involves both Th1 and Th2 activity and intestinal IL-10 balances these responses, promoting Th2 cytokine expression by limiting local Th1 cells.

The original identification of IL-10 was as an effector cytokine of Th2 immunity^18^, but there have been mixed reports of its regulatory impact on Th2 cells. He, Poholek and colleagues recently argued that IL-10 signalling is critical for the development of the Th2 response in a murine model of asthma^22^, and Coomes, Wilson and colleagues have previously reported that IL-10 promotes full Th2 differentiation in allergic airway inflammation^23^. In contrast, others have reported that IL-10 inhibits Th2 activity in the lung^23, 58^. Even within a single parasite infection with *Trichinella spiralis*, IL-10 can both suppress Th2 immunity in infected muscle and promote Th2 activity in infected instestine^25^. Site-specific variation in IL-10 signalling complements our growing understanding of the importance of tissue-specific regulation of immunity^59, 60^. One of the location-dependent factors that could influence IL-10’s impact might be the presence or absence of an underlying Th1 response. Despite containing fewer bacteria than the colon, the small intestine still has an abundant microbiome^61^ and any breach of the intestinal epithelium provides an opportunity for bacterial translocation and the stimulation of anti-bacterial immunity. In *H. polygyrus* infection, larvae and adult worms burrow into and out of the intestinal wall of the small intestine at days 1-2 and 7-8 of infection. Barrier disruption in the presence of intestinal bacteria has been hypothesised to lead to IFNγ expression^62^, and our data showing Th1 expansion in the *H. polygyrus*-infected intestine when IL-10 is blocked provide new experimental evidence. Bacterial translocation has been reported in other infections that damage the integrity of the intestinal wall, such as in *Toxoplasma gondii*, where the microbiota-specific T cell response has been shown to amplify the parasite-specific Th1 response to infection^63^. In helminth infection, where protective immunity is Th2 biased, a Th1 component to the anti-parasite immune response may instead act as a competitive inhibitor^39^.

*H. polygyrus* infection is also associated with a shift in the balance of bacterial species in the intestine, favouring an expansion of Lactobacillae^64^, and this altered microbiome might alone be immunogenic ^62, 64, 65^. In *Trichuris muris* infection, Duque-Correa, Berriman and colleagues have reported that IL-10 can influence both the composition of the intestinal microbiome and the translocation of those bacteria across the intestinal wall^66^. Here we report that IL-10 in *H. polygyrus* infection regulates an intestinal Th1 response, and it will be interesting in future studies to assess whether the underlying Th1 response is a reaction to helminth- and/or bacterial-derived stimuli.

In many protozoan and bacterial infections, the role of IL-10 in limiting IFNγ expression is critical for host survival, suppressing damaging immunopathology^33, 34, 67, 68^. IL-10 has also been proposed to limit tissue pathology in the intestine by promoting epithelial cell proliferation and subsequent colonic wound repair, via WISP-1 signalling^69^. Despite these data, we did not see significant changes in intestinal pathology in *H. polygyrus* infection when IL-10 signalling was blocked. Pathology in *H. polygyrus* infection is both mild and patchy, concentrated around the granulomas that encase developing larvae^10, 70^. A more detailed analysis of granulomas, or of later times of infection, may have revealed more marked differences, However, even at this early time point, our data did reveal striking differences in intestinal cytokine profiles. These data suggest that, in *H. polygyrus* infection, the main impact of IL-10 is to suppress IFNγ and to enhance Th2 function. The ability of IL-10 to promote the Th2 response was observed only in the small intestine lamina propria and not in the draining lymph node, complementing a growing understanding of the importance of tissue-specific regulation of immunity^59, 60^.

Immune regulation through the balance of opposing T cell cytokines is a common feature of infection. In mice, *H. polygyrus* infection of MyD88^−/−^ animals leads to reduced IFNγ expression, heightened IL-4, and accelerated parasite expulsion^71^. Mice without a functional IL-4 receptor show exaggerated IFNγ recall responses during *H. polygyrus* infection, compared with wildtype controls^39^. Cytokine exclusion is often less absolute in humans, but a recent report of a child with an inherited Tbet deficiency described elevated Th2 cytokine production^72^. The mechanisms of regulation by opposing Th1 and Th2 cytokines can include direct molecular inhibition of signalling within the CD4^+^ T cell, such as STAT1-driven induction of Tbet, and Tbet mediated suppression of GATA3^73^. Cytokine competition can also be achieved through different conditioning of antigen presenting cells, recruitment of different effector cells, or alteration of metabolic profiles^74^. Our data emphasise that such cytokine competition is a key feature of the intestinal immune response during enteric helminth infection, and that IL-10 is a key regulator of this process.

Our observation of high expression of the IL-10 receptor on Th1 cells in the intestine, greater than on intestinal Th2 cells or on all T cells in the draining lymph node, suggested strong, local IL-10 signalling to Th1 cells. Surface expression of cytokine receptors reflects both gene expression and surface binding, internalisation and recycling^75^ and in vitro, when exogenous cytokines are added at supraphysiological concentrations, active signalling can result in loss of surface expression of the receptors^76, 77^. *in vivo*, active cytokine concentrations are lower and cell surface receptor stripping is less commonly observed^78^; instead, increased receptor expression is associated with increased signalling^48, 79^. Macrophage expression of IL-10R1 determines their responsiveness to exogenous IL-10^49^. Decreased IL-10R1 expression on peripheral T cells in lupus patients correlates with heightened T cell activity, suggesting that reduced receptor expression is associated with reduced IL-10 function^80^. Our data show high expression of IL-10R1 on Th1 cells in the intestine, and increased frequency of intestinal Tbet^+^ or IFNγ^+^ T cells when IL-10 signalling is blocked. Together these data suggest that understanding receptor expression will be an important step in targeting attempts to use IL-10 therapeutically, which has been challenging and highly context-dependent^34, 81^.

In summary, we have shown that IL-10 promotes the intestinal Th2 response to a helminth infection. We show that IL-10 can signal directly to CD4^+^ T cells to increase Th2 differentiation, but that IL-10 production and IL-10 receptor expression are both concentrated in the infected tissue rather than the MLN. High expression of the IL-10R by Th1 cells in the small intestine suggests that these cells are sensitive to IL-10 signalling and subsequent IL-10 mediated suppression, providing an indirect mechanism in which IL-10 promotes the intestinal Th2 response. Our data provide new insight into the complexity of tissue-based regulation during a Th2 immune response and suggest that IL-10 may be an interesting candidate for therapeutic targeting in Th2 dominated diseases such as allergy, asthma and helminth infection.

## 4 Methods

### Mice and Infection

C57BL/6 mice were purchased from Envigo (Huntingdon, UK). B6.4get mice were kindly provided by Professor Judi Allen (University of Manchester) and bred in-house at the University of Glasgow. These mice were first developed by Mohrs *et al*^50^. Il10gfp-foxp3rfp B6 mice were bred in-house (University of Glasgow). These mice express two separate transgenes: IRES-eGFP inserted at the last exon and before the polyadenylation site of the Il10 gene^44^ and IRES-RFP inserted at this site of the Foxp3 gene^45^. For each experiment mice were sex-matched and used at age 6-12 weeks. Animals were maintained in individually ventilated cages under standard animal house conditions at the University of Glasgow and procedures were performed under a UK Home Office licence (Project number 70/8483) in accordance with UK Home Office regulations following review by the University of Glasgow Ethics Committee. Mice were acclimatised for 1 week after arrival in the animal unit before use. For infections, *H. polygyrus* (also known as *H. polygyrus bakeri* or *H. bakeri*) was maintained in the laboratory as described^82^, and experimental animals were infected with 200 L3 larvae by oral gavage.

### Isolation of cells

Lamina propria leukocytes were isolated as described previously^40^. The MLN was harvested and crushed through a 70μm filter to obtain a single cell suspension. For experiments where myeloid cells were analysed, MLNs were digested for 40min in a shaking incubator using 1mg/ml collagenase D (Merck) in RPMI. Cells were counted and dead cell exclusion carried out using trypan blue.

### *In vitro* CD4^+^ T cell culture and proliferation

Negative selection of CD4^+^ T cells from naïve splenocytes was carried out using the MojoSort™ magnetic cell separation system (Biolegend). CD4^+^ T cells were re-suspended in in RPMI 1640 supplemented with 10% FCS, 100 U/ml penicillin, 100μg/ml streptomycin and 2mM L-glutamine and stimulated in a 96-well plate with plate bound αCD3 (1μg/ml), soluble αCD28 (1μg/ml) and appropriate stimulation and polarisation cocktails. Polarisation cocktails: Th0: 20ng/ml IL-2, Th2: IL-2 (20ng/m), IL-4 (40ng/ml) (ThermoFisher), αIFNγ (1μg/ml) (Biolegend). Th1: IL-2 (20ng/m), IL-12 (10ng/ml) (ThermoFisher). Cells were then cultured for 4 days 37°C, 5% CO_2_. For IL-10 stimulation, IL-10 (ThermoFisher) was added at 10ng/ml. For assessing CD4 T cell proliferation the CellTrace™ Violet Cell Proliferation Kit (ThermoFisher) was used according to manufacturer’s guidelines.

### Ex-vivo re-stimulation

To measure cytokine production following ex vivo re-stimulation, unfractionated MLN cells were resuspended at 5×10^6^ cells/ml in RPMI 1640 supplemented with 10% FCS, 100 U/ml penicillin, 100μg/ml streptomycin and 2mM L-glutamine. 500,000 cells were then added to each well, coated with αCD3 (1μg/ml). Cultures were incubated (37°C, 5% CO2) for 3 days and then supernatants collected for further analysis.

### Cytokine measurement in supernatants

Supernatants were collected from *in vitro* T cell cultures or ex-vivo stimulated cultures and stored at −20°C until further analysis. For cytokine measurements, supernatants were diluted 1/200 in sterile filtered FACS buffer (PBS, 2mM EDTA and 10% FCS). Cytokines (IL-5 and IL-13) were measured using BD™ CBA Flex Sets (BD Biosciences) according to manufacturer’s guidelines. The cytometric bead array was analysed using the MACSQuant® Analyser (Miltenyi Biotec). Analysis was performed using FlowJo (Treestar).

### Flow cytometry and intracellular cytokine staining

To measure cytokine production immediately ex vivo, cells were stimulated and then stained for flow cytometry. 3×10^6^ cells were resuspended in in 500μl of RPMI 1640 supplemented with 10% FCS, 100 U/ml penicillin, 100μg/ml streptomycin, 2mM L-glutamine and 2μl/ml solution of cell stimulation cocktail and protein transport inhibitors (Invitrogen eBioscience™ Cell Stimulation Cocktail plus protein transport inhibitors (500X)). After 4 hours of stimulation, cells were washed and stained with Fixable viability dye eFluor 780 or 506 (Ebioscience), to enable dead cell exclusion, and then with anti-mouse CD16/32 Antibody (BioLegend) as an FC block, to reduce non-specific binding. Samples were next stained for surface markers for 20min at 4°C with: PerCP-Cy5.5-conjugated anti-TCRβ (H57–597, BioLegend), APC-CY7-conjugated anti-B220 (RA3-6B2, BioLegend), APC-CY7-conjugated MHCII (M5/114.15.2, eBioscience), APC-conjugated anti-IL-7Rα (A7R34, BioLegend), BV421-conjugated anti-CD44 (IM7, BioLegend), PE-Cy7-conjugated anti-CXCR3 (CXCR3-173, BioLegend), PE-conjugated IL-10R (1B1.3a, BioLegend), APC-Cy7-conjugated anti-CD19 (6D5, BioLegend) BUV395-conjugated CD45 (30-F11, BD Bioscience), BV711-conjugated anti-CD4 (RM4-5, BioLegend), BV421-conjugated anti-CD11b (M1/70, BioLegend), PE-Cy7-conjugated anti-CD8 (53.6.7, BioLegend), FITC-conjugated anti-CD44 (IM7, BioLegend) BV505-conjugated anti-CD4 (RM4-5, BioLegend), FITC-conjugated anti-CD69 (H1.2F3, BioLegend). Samples were then permeabilised and fixed for intracellular cytokine staining using 150μl of BD Cytofix/Cytoperm™ for 20 min at 4°C. Samples were stained using 50μl of intracellular anti-cytokine antibody stain: PE-Cy7-conjugated anti-IL-13 (eBio13A, Invitrogen), PE-conjugated anti-IL-5 (TRFK5, BioLegend), e450-conjugated anti-IFNγ (XMG1.2, Invitrogen)) or appropriate isotype controls for 1 hour at room temperature in the dark). When including staining for intracellular transcription factors, samples were then permeabilised and fixed intracellularly using the eBioscience™ Foxp3 / Transcription Factor Staining Kit (ThermoFisher) for 1 hour at room temperature in the dark. Samples were stained with 100μl of intracellular anti-transcription factor stain: eFluor 450-conjugated anti-FOXP3 (FJK-16s, ThermoFisher), PE-Cy7-conjugated anti-T-bet (eBio4B10, ThermoFisher) and PE-conjugated anti-GATA3 (TWAJ, ThermoFisher) for 1.5 hours at room temperature in the dark. All samples were acquired immediately after staining on either a BD LSRII flow cytometer or a BD LSR FORTESSA running FACS-Diva software (BD Biosciences). Analysis was performed using FlowJo (Treestar).

### RNA extraction and real-time PCR

1cm section of the top of the duodenum was collected, placed in RNA later (Qiagen) and kept at 4°C RNA was purified using RNEASY Mini Kit (Qiagen) and its concentration determined using a Nanodrop 1000. cDNA was generated using the High Capacity cDNA Reverse Transcription Kit (Invetrogen). For real time PCR, PowerUp™ SYBR™ Green Master Mix (Applied Biosystems) and QuantStudio 6 Flex Real-time PCR system (Applied Biosystems) were used. Values were normalised to ribosomal protein S29 (RPS29) and expression of gene of interest determined using the 2^−ΔΔC(t)^ method. The following primers were used (all primers are shown 5’-3’); *Rps*29 Fwd, ACGGTCTGATCCGCAAATAC, Rev, CATGATCGGTTCCACTTGGT; *Il*13 Fwd, CCTGGCTCTTGCTTGCCTT, Rev, GGTCTTGTGTGATGTTGCTCA; *Il*5 Fwd, CTCTGTTGACAAGCAATGAGACG, Rev, TCTTCAGTATGTCTAGCCCCTG; *Il*22 Fwd, TTTCCTGACCAAACTCAGCA, Rev, CTGGATGTTCTCGTCGTCAC.

### IL-10R1 monoclonal antibody blockade

IL-10R signalling was blocked using an IL-10R1 monoclonal antibody (Clone 1b1.3a BioXcell). A rat IgG1 antibody (Merck) was used as the isotype-matched control. 200μl of a 2.5mg/ml stock solution of each antibody was injected intraperitoneally at days −1, 2, and 5 of *H. polygyrus* infection. Infected mice treated with anti-IL-10R or isotype were kept in mixed cages.

### Histology and scoring

The first 6cm of the duodenum were collected, sliced into 1cm pieces and placed in 10% neutral buffered formalin. Samples were fixed overnight, trimmed and embedded in paraffin wax. Tissue sections were collected on frosted glass slides and stained with haematoxylin and eosin. The depth of inflammation, and a combined score of depth and severity of inflammation, were evaluated blindly by a certified pathologist (VG) in several high-power fields of two intestinal sections per animal using a protocol established to previously^83^. The scoring system was from 0-4: 0 – Minimal and mucosal, 1 – Mild and mucosal/submucosal or minimal transmural, 2 – Moderate and mucosal/submucosal, mild transmural or marked and mucosal, 3 – Marked mucosal/submucosal or moderate and transmural, 4-Marked and transmural.

### Statistical analysis

All statistical analysis was carried out using GraphPad Prism (version 8/9) and data represents mean + standard deviation. A Student t test was used for comparison between 2 groups and a one-way ANOVA with Tukey’s multiple comparison correction was carried out for comparisons between 3 or more groups. All data sets were tested for normality using the Shapiro-Wilk normality test and where data were not normally distributed, a Mann Whitney U test for comparisons between 2 groups and a Kruskal-Wallis test with Dunn’s multiple comparison correction was carried out for comparisons between 3 or more groups. \**p*<0.05, \*\**p*<0.01, \*\*\**p*<0.001, \*\*\*\**p*<0.0001, ns = not significant.

## 5 Acknowledgements

We thank David Dow, Joanne Battersby and staff in the Wolfson Research Unit for animal husbandry; Nicola Britton and Claire Ciancia for *H. polygyrus* larvae and life cycle maintenance, and for advice and discussion; Allan Mowat for critical review of the manuscript; the Flow Core at the University of Glasgow and especially Diane Vaughan of the manuscript for flow cytometry support; and Lucy McShane and Lotus Westerhof for sample processing. This work was supported by the MRC (grant MR/S009779/1) and funds from the University of Glasgow, including a University of Glasgow MVLS PhD studentship awarded to HCW. RMM also received Wellcome Trust support through an Investigator Grant (Ref 219530), and the Wellcome Trust core-funded Wellcome Centre for Integrative Parasitology (Ref: 104111).

## 6 Author Contributions

H.C.W conceived and refined the experimental and conceptual design of the study, conducted experiments, analysed data and prepared the manuscript. V.G is a veterinary pathologist and conducted all analysis of the histopathology and developed a scoring system for histological samples. A.L.S, A.T.A and G.A.H performed experiments and acquired data. S.W.F.M provided critical expertise and edited the manuscript. R.M.M contributed to the conceptual design of the study, provided critical expertise and edited the manuscript. G.P.W conceived and refined the experimental and conceptual design of the study, analysed data and prepared and edited the manuscript.

## 7 Disclosures

The authors declare no competing interests.

## Notes

**Disclosures:** All authors state that they have no conflicts of interest in the content of this manuscript.

### Competing Interest Statement

The authors have declared no competing interest.

