## Supplemental data for "Tissue-based IL-10 signalling in helminth infection limits IFNγ expression and promotes the intestinal Th2 response"

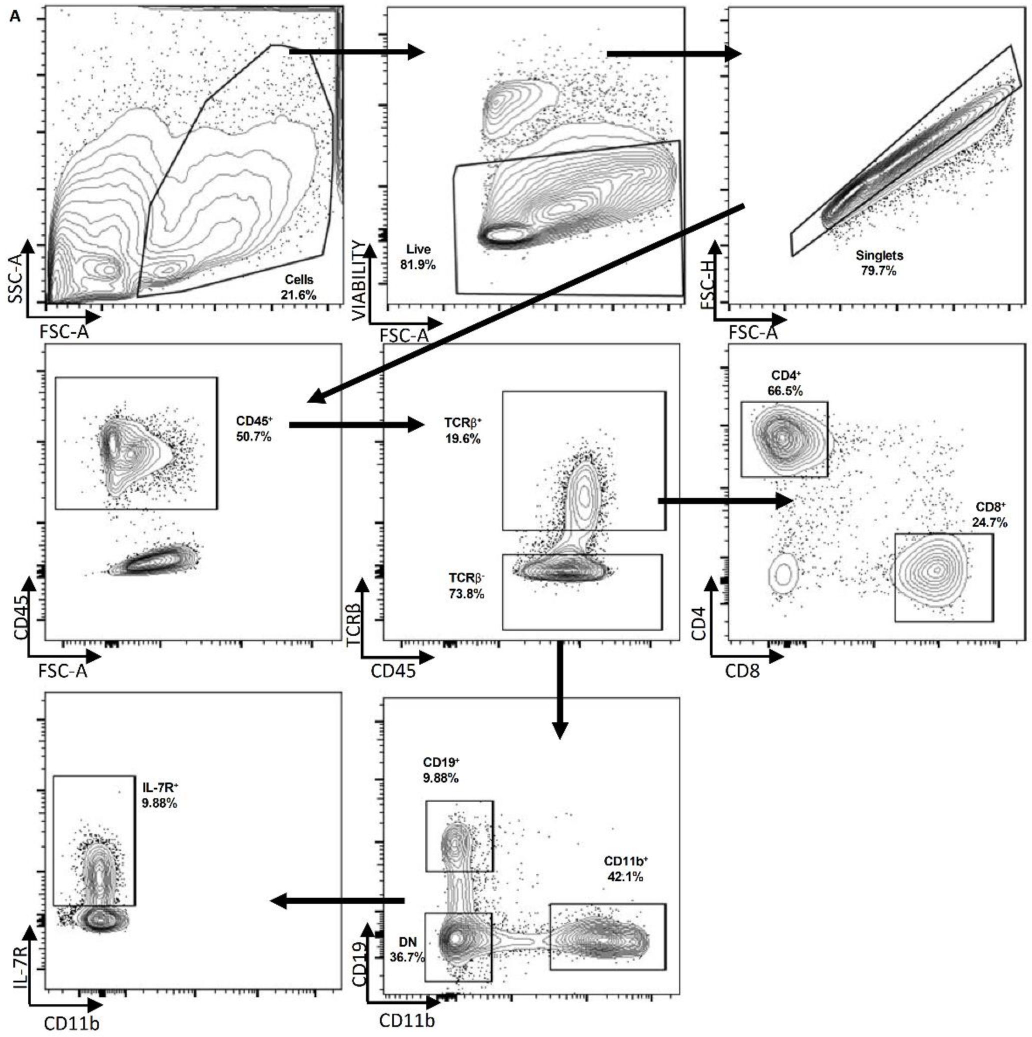


**Supplementary Figure 1 Gating for cell subsets in the small intestine during H. polygyrus infection**

Il10gfp-foxp3rfp B6 mice were infected with 200 L3 *H. polygyrus* and 7 days later the small intestine and MLN collect for analysis. Representative gating of cell subsets from D7 *H. polygyrus* infected small intestine.

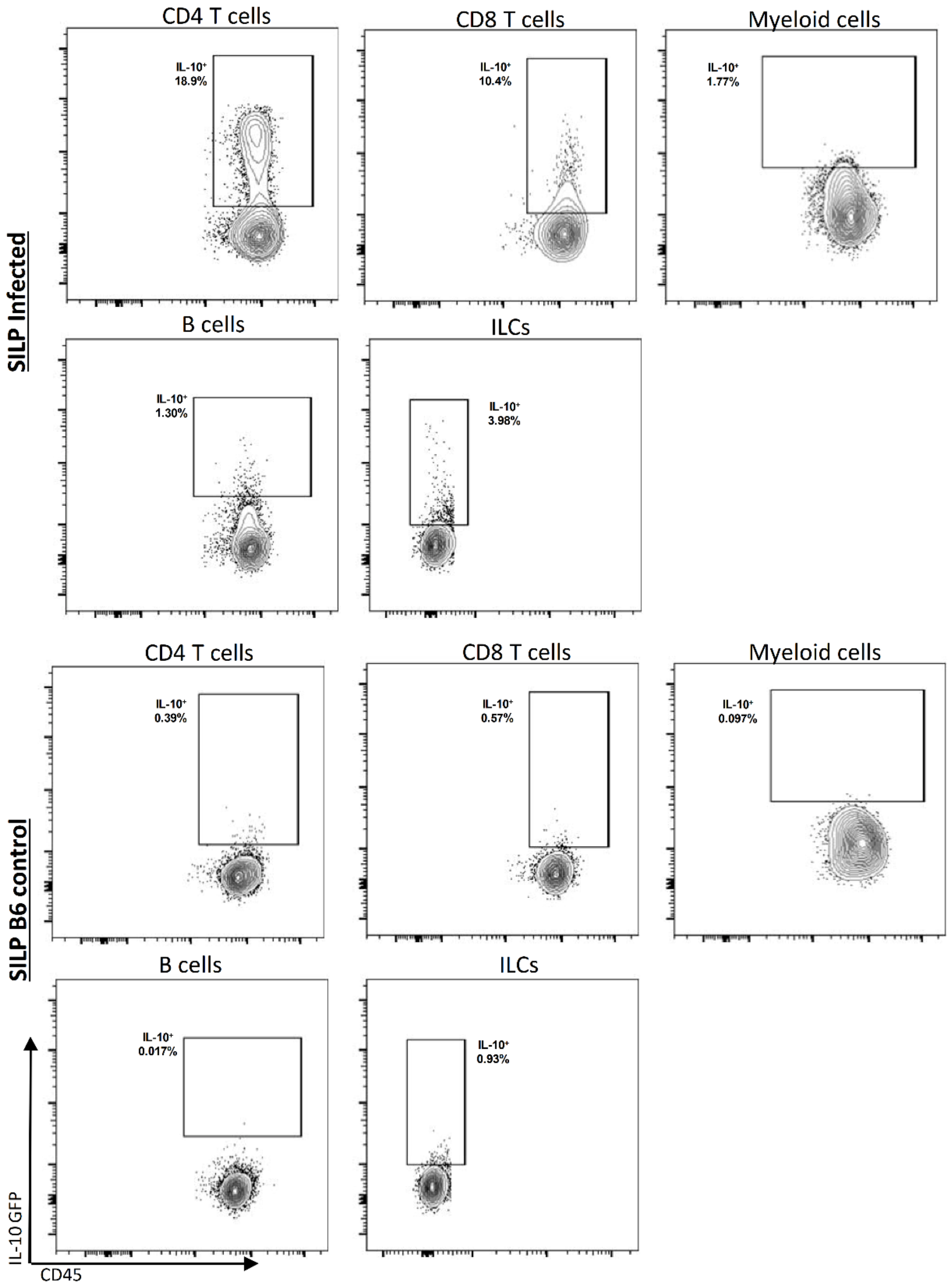


**Supplementary Figure 2 IL-10 (GFP) gating for cell subsets**

Il10gfp-foxp3rfp or C57BL/6 mice were infected with 200 L3 *H. polygyrus* and 7 days later the small intestine and MLN collected for analysis. Representative gating for IL-10 from each cell subset from D7 *H. polygyrus* infected small intestine and B6 control mice. All gates were drawn using B6 control mouse as a negative control.

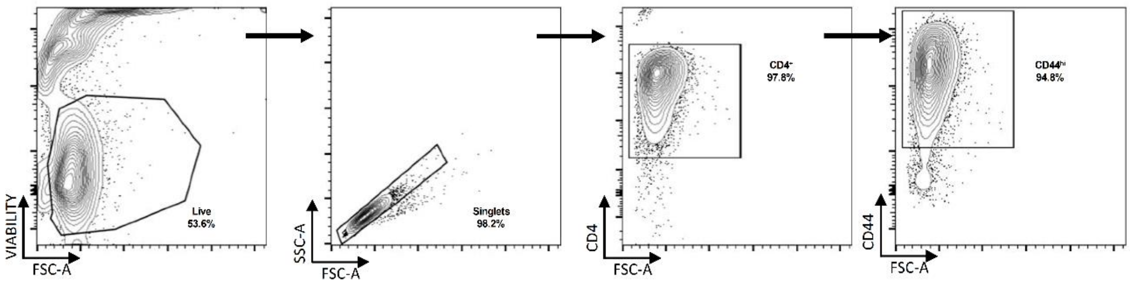


**Supplementary Figure 3 In vitro CD4 T cell gating**

*In vitro* polarised Th0 cells were cultured with αCD3, αCD28 and IL-2 for 4 days and harvested for further analysis. Representative gating strategy of *in vitro* activated Th0 cells.

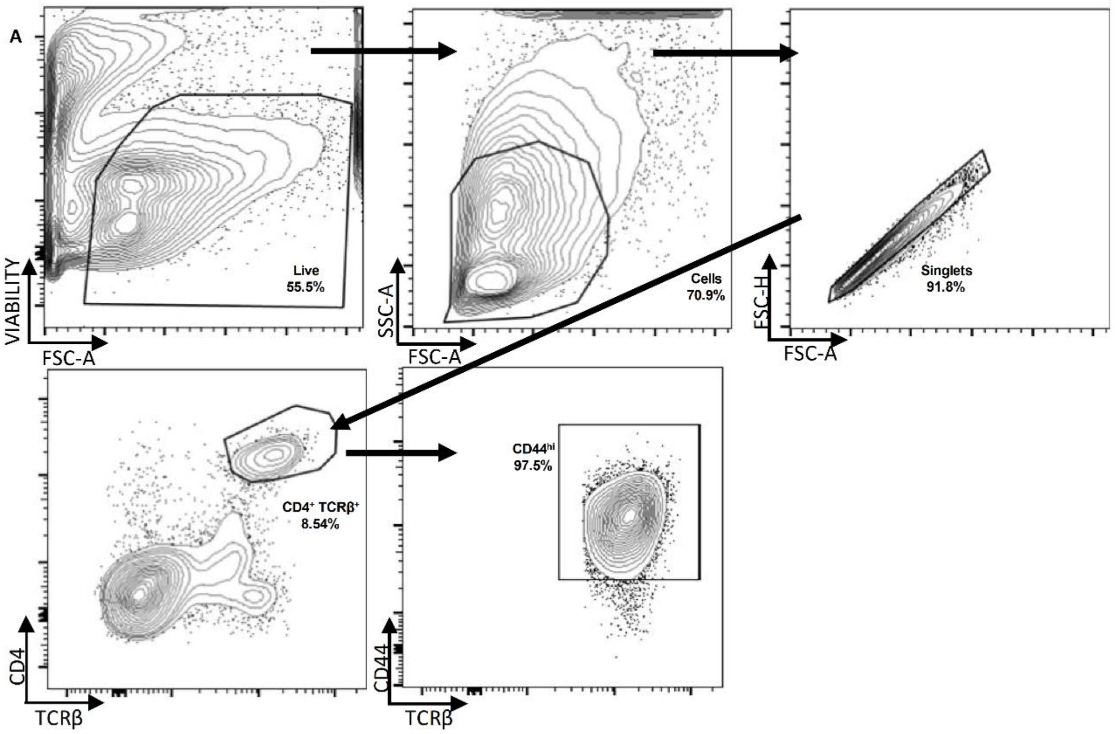


**Supplementary Figure 4 In vivo CD4 T cell gating**

C57BL/6 mice were infected with 200 L3 *H. polygyrus* and at D-1, D2 and D5 treated with anti-IL-10R mAb or isotype control,  and 7 days post-infection the small intestine and MLN collect for analysis. Representative gating of activated CD4 T cells from D7 *H. polygyrus* infected small intestine.

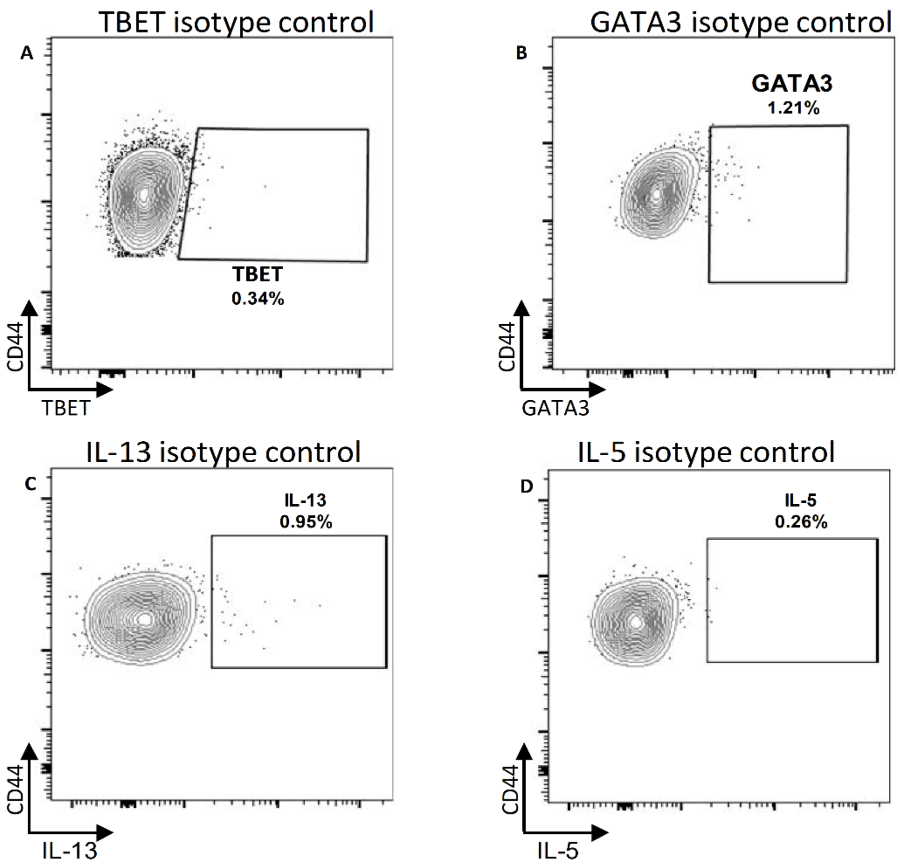


**Supplementary Figure 5 Isotype controls for ex vivo intracellular staining of CD4 T cells**

C57BL/6 mice were infected with 200 L3 *H. polygyrus* and at D-1, D2 and D5 treated with anti-IL-10R mAb or isotype control,  and 7 days post-infection the small intestine and MLN collect for analysis. Representative gating using intracellular isotype controls for (A) TBET, (B) GATA3, (C) IL-13 and (D) IL-5 cells from CD4$^+$ T cells from D7 *H. polygyrus* infected small intestine

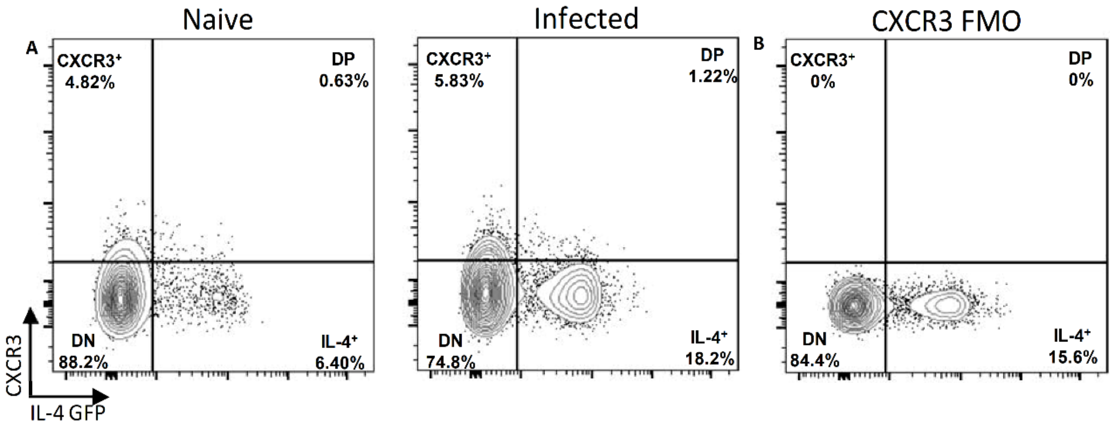


**Supplementary Figure 6 IL-4 and CXCR3 staining**

B6 4get mice were infected with 200 L3 *H. polygyrus* and 7 days post-infection the small intestine and MLN removed. (A) Representative staining for IL-4 and CXCR3 expression by CD4 T cells from D7

*H. polygyrus* naïve and infected small intestine.  (B) Representative staining from CXCR3 FMO control used to gate on CXCR3 positive cells.
